# Domestication and Improvement Reshaped the Genomic Architecture of Abiotic Stress Tolerance in Sunflower

**DOI:** 10.64898/2025.12.03.692181

**Authors:** Yavuz Delen, Semra Palali-Delen, Mahamkali VS Sai Subhash, Haichuan Wang, Ismail Dweikat, Jinliang Yang, Gen Xu

## Abstract

Sunflower (*Helianthus annuus*) has undergone extensive domestication and modern improvement, yet the extent to which selection has shaped its stress-adaptive variation remains largely unclear. Using genomic data from diverse accessions, including the wild ancestor of sunflower, we identified widespread balancing selection that maintains historical polymorphisms in abiotic-stress-related genes. Additionally, we identified strong positive selection signals associated with domestication and improvement. Using a cutting-edge DNA language model to characterize deleterious alleles, we found that modern cultivars exhibited an elevated deleterious mutation load, particularly within positively selected regions, consistent with patterns observed in other major domesticates. To cross-validate the functional consequences of the selected regions on stress responses, we assessed salt tolerance in an independent association panel and evaluated ancestral allele effects on salt-responsive traits. Although only a small subset of the selected regions reached genome-wide significance, several ancestral alleles within positively selected sweeps tended to enhance salt tolerance. Together, these results indicate that recent selection may have unintentionally eliminated historically beneficial alleles, reshaping the genomic landscape of sunflower stress responses. This study highlights a breeding strategy to recover ancestral beneficial alleles for enhancing abiotic stress tolerance in modern cultivars.

## Introduction

Sunflower (*Helianthus annuus L*.) was domesticated in North America about 4,000 years ago, according to archeological, morphological, and recent genetic evidence (Harter et al., 2004; Heiser, 1951; Smith, 1989). Today, it is widely cultivated as a major oilseed crop and used for various purposes, including snack foods, animal feed, and industrial applications (Adeleke and Babalola, 2020). As a staple oil source in many countries, seed oil content and quality have been key targets of selection throughout sunflower improvement and remain central goals of modern breeding programs (Badouin et al., 2017).

Sunflowers are highly adaptable and can thrive in environments where many other crops fail, making them particularly valuable in regions affected by soil salinity (Shi and Sheng, 2005). Its robust root system and inherent tolerance to multiple abiotic stresses enable growth across diverse ecological zones, including areas with high salt levels (Rauf et al., 2012). Soil salinity, affecting roughly 7% of the world’s cultivable land, poses major challenges for crop establishment and yield, especially in arid and semi-arid regions where sunflower is commonly grown (Kang et al., 1996). High salinity disrupts germination, seedling emergence, water uptake, and cellular metabolism, leading to stunted growth, leaf chlorosis, and significant reductions in both seed and oil yield (Pessarakli, 2019; Xiong and Zhu, 2002). Given its natural resilience, sunflower represents a promising salt-tolerant crop for sustaining production in increasingly saline and climate-stressed agricultural landscapes (Wahid et al., 1999).

Like many domesticated crops (Huang et al., 2012; Hufford et al., 2012; Lin et al., 2014; Qi et al., 2013), sunflowers experienced a substantial genetic bottleneck during domestication (Tang and Knapp, 2003), and modern breeding efforts that focused heavily on improving oil content have likely overlooked valuable salt-tolerance alleles. Although sunflower is considered moderately salt-tolerant, substantial variation among germplasm underscores the need for improved, efficient screening approaches (Chen et al., 2024; Li et al., 2020). With the rapid expansion of genomic resources in sunflowers (Bercovich et al., 2022; Huang et al., 2023; Yi et al., 2025), there is now a renewed opportunity to recover naturally occurring salt-tolerant or other stress-resilient alleles from diverse cultivars, native American landraces, and wild relatives.

In this study, we leverage publicly available genomic resources that span wild, landrace, and modern cultivated sunflower accessions to perform a comprehensive population genomic analysis. Through these analyses, we identify historically balanced loci that are preliminarily associated with abiotic stress responses, including putative salt-tolerance sites. Within these regions, we also observed enrichment for deleterious variants, and our analyses suggested that some beneficial ancestral alleles may have been lost during domestication and modern improvement. Together, these results underscore the need for targeted breeding efforts to restore historically advantageous alleles related to salt tolerance. By providing a clearer picture of salt-related genetic variation and its evolutionary history, this study offers valuable guidance for developing more resilient sunflower cultivars.

## Materials and Methods

### Genetic materials and variant calling procedure

We downloaded raw sequence data from 332 sunflower accessions, including modern cultivars (*n* = 288), landraces (*n* = 18), and wild relatives (*n* = 26), from a previous study (Hübner et al., 2019). We then processed raw sequencing reads with fastp (version 0.20.0) to trim adapter sequences and remove low-quality bases before downstream analyses (Chen et al., 2018). Next, the cleaned reads were aligned to the reference genome HanXRQr2.0-SUNRISE (Badouin et al., 2017) using Bowtie2 (version 2.4.4) (Langmead and Salzberg, 2012). Subsequently, we removed the duplicated reads using Picard tools (version 2.18) and conducted SNP calling using Genome Analysis Toolkit’s (GATK, version 4.1) HaplotypeCaller (McKenna et al., 2010), with the following parameters: QD *<* 2.0, FS *>* 60.0, MQ *<* 20.0, MQRankSum *<* −12.5, and ReadPosRankSum *<* −8.0, to filter out low-quality SNPs. In addition, we removed SNPs with missing rates *>* 50%.

### Principal component analysis (PCA) and phylogenetic tree

We performed PCA using SNP dataset with plink1.9 (Chang et al., 2015). Using the same software, we computed identity-by-state (IBS) values among the 332 sunflower accessions with the parameter “–ibs-matrix”. The resulting IBS similarity matrix was converted into a genetic distance matrix (1 − IBS), which was then used to construct a neighbor-joining (NJ) phylogenetic tree using the ape package (Paradis and Schliep, 2019) in R (v4.3.0). The tree was visualized with a circular layout using the “ggtree” package (Yu et al., 2017).

### Salt treatment experiment at the seedling stage

We evaluated salt tolerance at the seedling stage using an independent association panel of 274 accessions (Delen et al., 2024). All accessions used in this experiment came from the United States Department of Agriculture, Agricultural Research Service (USDA-ARS), North Central Regional Introduction Station (NCRPIS) in Ames, Iowa, USA. For the experiment, we treated seeds with a fungicide and placed them in petri dishes. Under control conditions, we germinated the seeds in pure water, whereas under salt treatment, we germinated them in a 200 mM NaCl solution. We maintained all petri dishes in a 27 °C growth chamber for four days. After incubation, we recorded the germination ratio and radicle length for each accession, using three biological replicates per treatment.

We calculated salt-responsive (SR) traits using the following formula.

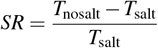

where *T*_nosalt_ and *T*_salt_ are the mean values for a given trait measured from non-salted and salted conditions.

### Genome-wide scan for positive and balancing selection

We performed a genome-wide scan for positive selection signals using the latest version of the cross-population composite likelihood ratio (XP-CLR) (Chen et al., 2010). In the XP-CLR analysis, we used a 50 kb sliding window and a 5 kb step size. To ensure comparability of the composite likelihood score in each window, we fixed the number assayed in each window to 100 SNPs. We then conducted a balancing selection scan with default parameters using BalLeRMix (version 2.3) (Cheng and DeGiorgio, 2020). Using the major alleles in wild species as ancestral alleles, we calculated the derived allele frequency (DAF). For both positive and balancing selection, we considered the 1% outliers as the selective sweeps; adjacent sweeps separated by a physical distance of *<* 50 kb were merged into a single selected sweep.

### Identification of deleterious mutations using a DNA language model

We used Plant Caduceus (Zhai et al., 2025), a DNA language model that infers the relative fitness effects of nucleotide substitutions through a zero-shot comparison of reference and alternate allele probabilities, to compute site-specific zero-shot scores (ZSS) and estimate genome-wide deleterious genetic load across wild, landrace, and improved sunflower groups. Variants were then classified into the top 1%, 5%, and 10% most deleterious categories based on their ZSS distributions, with more negative ZSS values indicating stronger predicted deleterious effects.

To test whether deleterious variants are enriched in genomic regions under selection, we intersected the genomic coordinates of deleterious SNPs with regions identified as under positive selection or balancing selection. For each category, we calculated the observed number of deleterious SNPs within selected regions and compared it with a null distribution generated from 1,000 sets of randomly sampled genomic windows matched in size.

### Genome-wide association analysis

We performed a genome-wide association analysis (GWAS) for salt responsiveness of germination ratio and radicle length using GEMMA (Zhou and Stephens, 2012), based on 226,779 high-quality SNPs generated in our previous study (Delen et al., 2024). We applied a mixed model Q + K to account for the confounding effects of both population structure (Q) and relatedness (K) (Yu et al., 2006). The Q matrix consisted of the first three principal components calculated using plink1.9 (Chang et al., 2015), while the K matrix was estimated with TASSEL 5.0 (Bradbury et al., 2007). To determine significant associations, we used a genome-wide threshold of 2.9 *×* 10^−5^, corresponding to 1*/n*, where *n* = 33,801 represents the number of independent SNPs. The number of independent markers was estimated using PLINK’s “indep-pairwise” function (window size = 10 kb, step size = 10, *r*^2^ ≥ 0.1). The significant GWAS loci were then defined by extending 50 kb upstream and downstream of each significant SNP, and the overlapping windows were merged into single genomic intervals. We estimated the sizes of allele effects using the beta coefficients from the GEMMA output. To determine the ancestral allele, we assigned the allele with the highest frequency in the wild group as the ancestral allele. A positive effect of the ancestral allele indicates that the ancestral allele contributes to a higher phenotypic value than the derived allele.

## Results

### Balancing selection maintains historical alleles associated with stress adaptation

To characterize the genetic relationships of the sunflower accessions, we first performed a principal component analysis (PCA) using 11.3 million genome-wide SNPs from 332 accessions (Hübner et al., 2019), including wild (*n* = 26), landrace (*n* = 18), and modern cultivated sunflower lines (*n* = 288) (**Table S1, Materials and Methods**). The PCA result clearly separated wild accessions from the cultivated group, with landraces positioned between them, with several outliers (**Figure 1A**). This pattern is largely consistent with a gradual transition during domestication and improvement processes observed for other domesticated crops (Hufford et al., 2012; Qiu et al., 2017; Wu et al., 2022; Zhou et al., 2015). The outlier accessions clustered closely with wild sunflower, likely reflecting ancestral genomic introgression as previously reported (Hübner et al., 2019).

**Figure 1.**
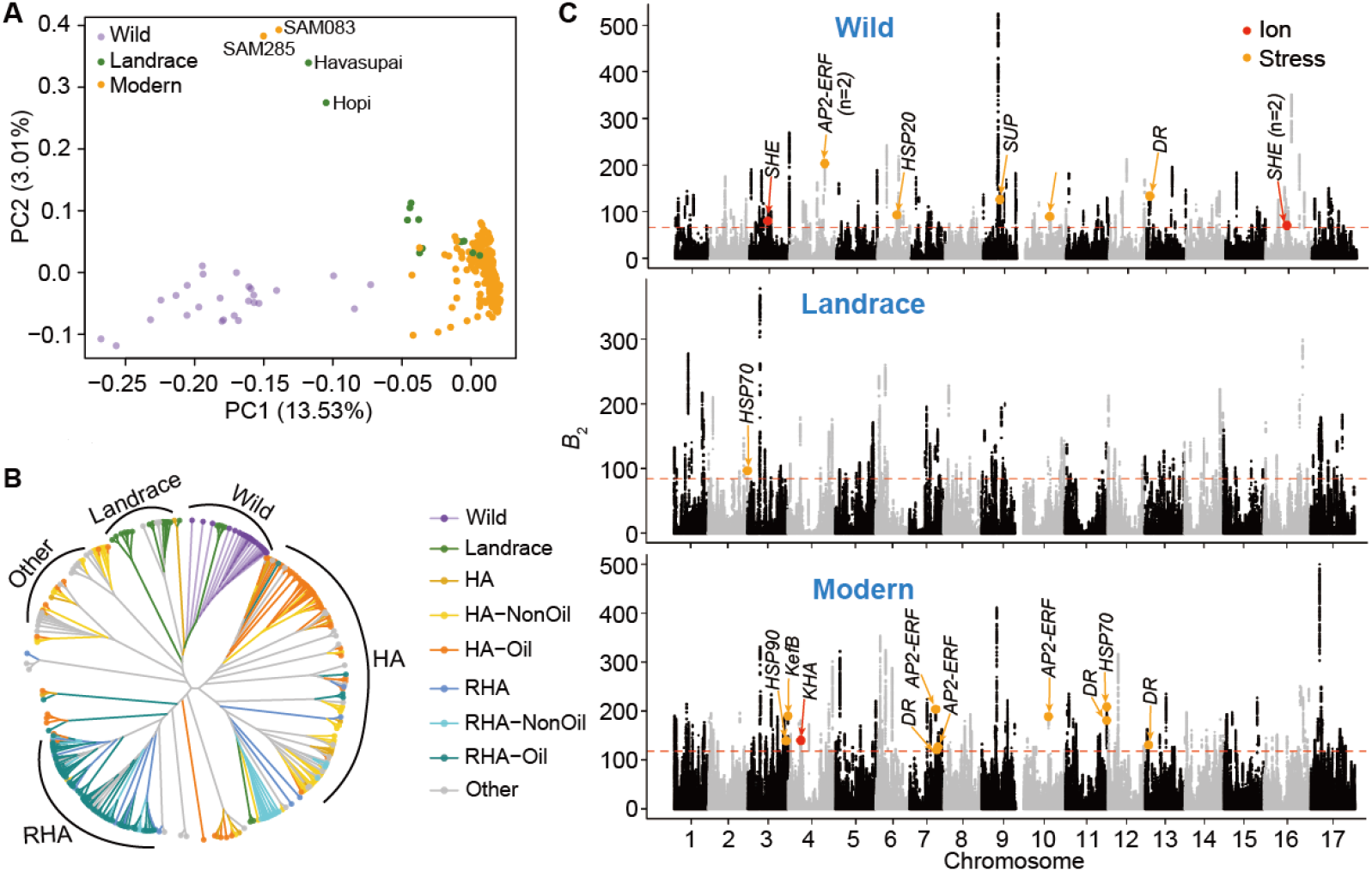
Genomic regions under balancing selection and their relationship with salt-stress-responsive loci in sunflower. (**A**) Principal component analysis (PCA) of the 332 accessions showed clear population differentiation among wild, landrace, and modern accessions. (**B**) Neighbor-joining phylogenetic tree of the modern accessions constructed from genome-wide SNPs. Colors represent different groups according to the previous study (Hübner et al., 2019). HA and RHA represent female and male lines, respectively. (**C**) Genome-wide balancing selection signals in each population, with colored dots indicating candidate genes involved in ion transport (red) and stress-responsive pathways (orange). The red horizontal dashed lines represent the significance level of the top 1%. SHE, sodium hydrogen exchanger; AP2-EREBP, ethylene-responsive transcription factor; HSP, heat shock protein; SUP, stress up-regulated; DR, disease resistance; KefB, potassium-efflux system protein; HKA, putative potassium channel.

We further investigated the genetic relationships among modern accessions by building a neighbor-joining phylogenetic tree (**Materials and Methods**). The tree showed a distinct clustering of female (HA) and male (RHA) lines in the cultivated group, corresponding to two heterotic groups used in hybrid breeding programs (**Figure 1B**). Within each group, moderate genetic diversity was observed, reflecting their different breeding origins and partial overlap between the oilseed and confectionery types, consistent with previous results (Mandel et al., 2011).

To explore how long-term evolutionary processes contribute to sunflower adaptive variation, we conducted a balancing selection scan and identified hundreds of genomic regions under balancing selection within each group (**Figure 1C, Table S2, Materials and Methods**). These balancing selection signals were widely distributed throughout the genome, overlapping with several candidate genes annotated as sodium–hydrogen exchangers (*n* = 3 in the wild population), heat shock proteins (*n* = 1 in the wild, *n* = 1 in the landrace and *n* = 2 in modern cultivars) and EREBP-type transcription factors (*n* = 3 in the wild and *n* = 3 in modern cultivars), which are associated with ion transport and stress-response pathways (**Table S3**). Such loci may represent allelic variants maintained by balancing selection that contribute to abiotic stress adaptation, such as salt tolerance.

### Positive selection drives genomic differentiation during historical domestication and recent improvement

Next, we investigate genomic footprints of positive selection during sunflower domestication and improvement processes (**Figure 2**). Using the cross-population composite likelihood ratio (XP-CLR) approach (**Materials and Methods**), we identified a total of 652 domestication-related selective sweeps (landraces vs. wild accessions) and 549 improvement-related sweeps (modern accessions vs. landraces), covering 2.3% (67 Mb) and 1.9% (56.9 Mb) of the genome, respectively (**Table S4**). Domestication sweeps ranged from 50 to 910 kb, with an average length of 103 kb and a total of 1,919 genes. During the improvement process, a slightly smaller number of regions was detected, with a comparable mean footprint size (104 kb), which encompassed 2,062 genes. The genes within these regions include key functional categories such as kernel oil biosynthesis (19 genes in domestication; 11 in improvement), flowering-time regulation (*n* = 6 in domestication; *n* = 13 in improvement), ion transport (*n* = 9 in domestication; *n* = 3 in improvement), and stress-response pathways (*n* = 18 in domestication; *n* = 20 in improvement, **Table S5**). The differences in category composition suggest that domestication disproportionately targeted loci associated with early agronomic traits, particularly oil accumulation and ion homeostasis. In contrast, modern improvement more strongly emphasized flowering regulation and stress-resistance functions. In particular, nine domestication sweeps showed evidence of continued selection during improvement (**Figure 2**), suggesting that a subset of key loci remained under strong selection pressure throughout the history of sunflower improvement.

**Figure 2.**
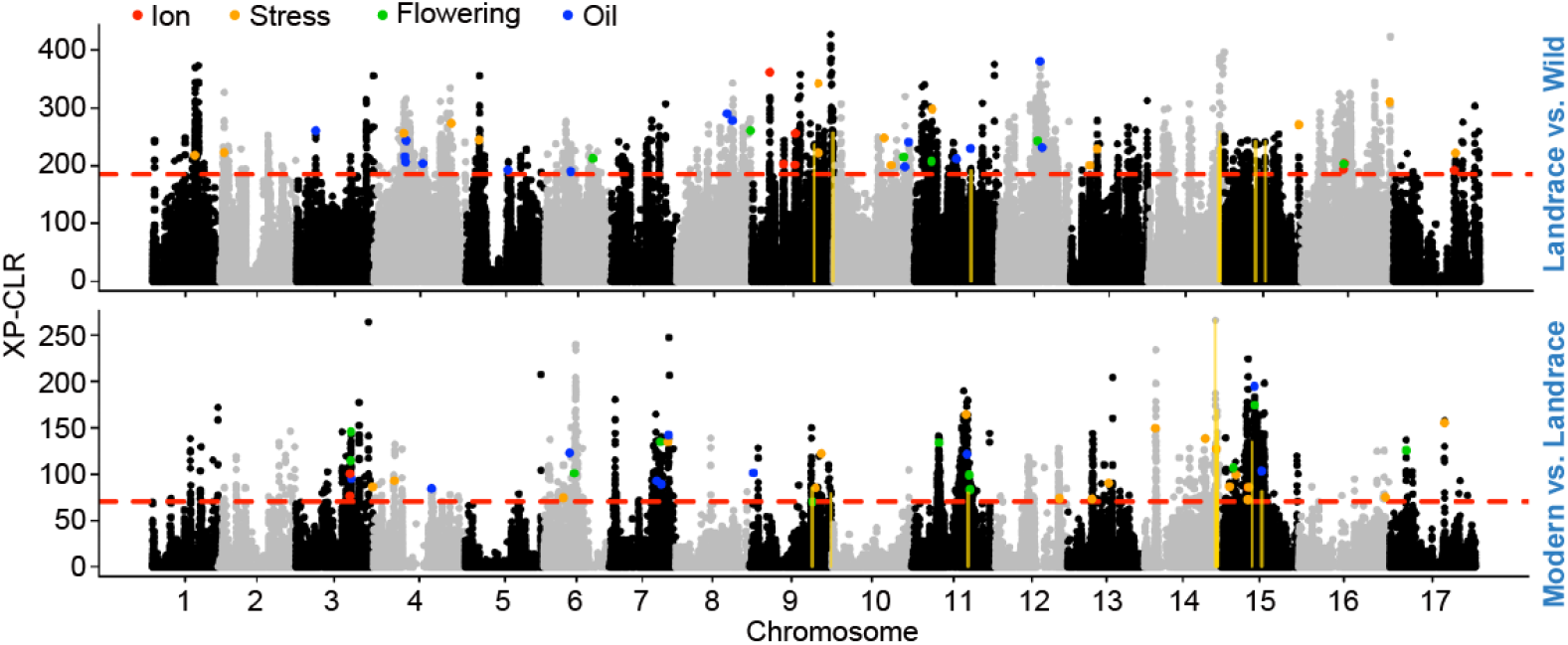
Genomic regions under positive selection during sunflower domestication and improvement. Genome-wide XP-CLR analysis was performed to detect genomic regions under differential selection between landraces and wild accessions (top) and between modern cultivars and landraces (bottom). The red dashed line indicates the top 1% threshold used to define candidate selective sweep regions. Annotated genes within these regions are categorized by their putative biological functions related to ion transport (red), stress response (orange), flowering regulation (green), and oil biosynthesis (blue). The gold vertical lines indicate the overlapped domestication and improvement sweeps.

### Domestication and selection processes shaped deleterious genetic load in sunflower

To assess the genome-wide burden (or genetic load) of deleterious mutations in sunflower, using a pretrained DNA language model (Zhai et al., 2025), we calculated zero-shot scores (ZSS) for 11.3 million SNPs (**Materials and Methods**). In the analysis, ZSS is calculated as the logarithmic probability of the alternative allele relative to the reference allele, with more negative values indicating a higher predicted deleteriousness as evidenced by the relationship between the frequency of the minor allele and the ZSS values (**Figure S1**). The ZSS values exhibited a broad distribution (**Figure 3A**). To identify deleterious sites, instead of relying on a single arbitrary threshold, we defined the upper-tail cutoff points of 1%, 5%, and 10% as increasingly relaxed thresholds (**Figure 3A**). Using these thresholds, although the total number of deleterious alleles varies, a consistent pattern emerged: modern cultivars carried the highest number of deleterious alleles, followed by landraces, whereas wild accessions harbored the lowest load (**Figure 3B**). This pattern is consistent with the increased accumulation of mildly deleterious mutations observed in other domesticated crops, such as maize (Lozano et al., 2021), wheat (Fu and Horbach, 2025), and rice (Lu et al., 2006). In addition, we evaluated the deleterious genetic load within regions under selection. Regions experiencing positive selection during both domestication and improvement showed significantly higher counts of deleterious SNPs than expected from randomly sampled genomic intervals (**Figure 3C**). This enrichment was consistent across all three deleteriousness thresholds, supporting the idea that selective sweeps may have carried linked deleterious alleles to higher frequencies. Similarly, regions under balancing selection also exhibited an enrichment of deleterious mutations, though to a lesser extent (**Figure 3D**), indicating that long-term maintenance of polymorphism may preserve some slightly deleterious variants.

**Figure 3.**
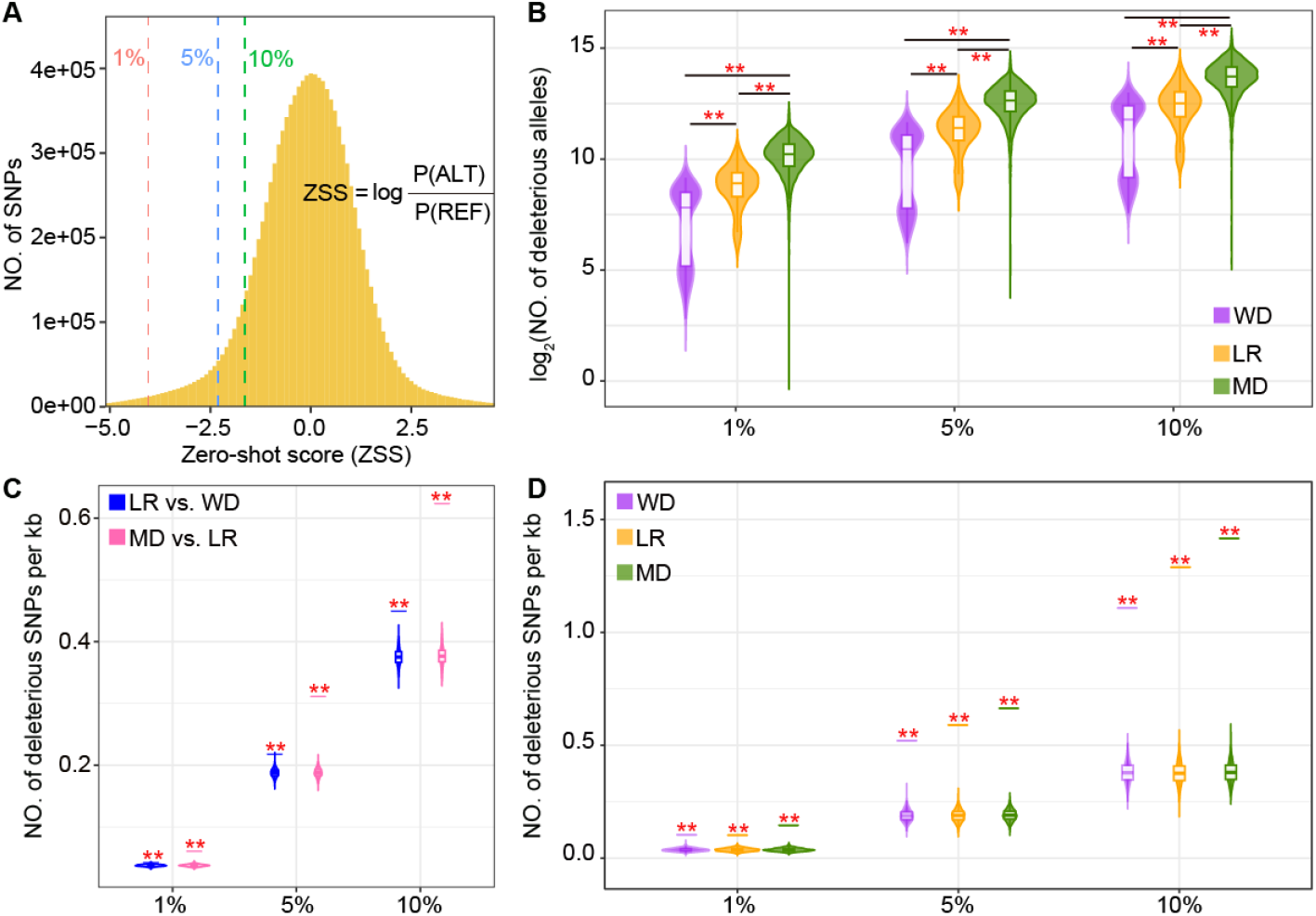
Identification of deleterious mutations in sunflower. (**A**) Zero-shot score (ZSS) distribution of 11.3 million SNPs in the sunflower population. The 1%, 5%, and 10% cutoffs indicate increasingly relaxed deleteriousness thresholds, with 1% representing the most deleterious variants. (**B**) Distribution of genetic load (sum of ZSS) across wild (WD), landrace (LR), and modern (MD) accessions under the three deleteriousness thresholds (1%, 5%, and 10%). (**C–D**) Deleterious mutations located within regions under positive (**C**) and balancing (**D**) selection. Violin plots show the distribution generated from 1,000 randomly selected genomic regions, and short horizontal lines indicate the observed values. Asterisks denote significance from permutation tests (**, *P* ≤ 0.01).

### Salt-response variation is shaped by selection and reflected in population-differentiated allele effects

Given that numerous genes related to stress, particularly salt-response, are located within regions under balancing or positive selection, we hypothesize that adaptive responses to abiotic stress have played an important role throughout the history of sunflower improvement. To directly connect these evolutionary signatures with functional phenotypes, we performed a salt-stress experiment using an independent association panel and quantified genotype-specific responses (**Materials and Methods**). As expected, both germination ratio (GR) and the radicle length (RL) decreased significantly with salt treatment (**Figure S2A-B**). Beyond raw traits, we also computed salt-responsive (SR) values following a previously described method (Liu et al., 2021; Palali Delen et al., 2023). In general, RL exhibited a stronger salt response than GR (**Figure S2C**) (Wilcoxon test, *P* = 5.2 *×* 10^−66^).

We next conducted GWAS for the transformed SR traits using a linear mixed model (see **Materials and Methods**). For GR, we identified 42 significant trait-associated loci (TALs), six (15%) of which were located within regions under positive selection. For RL, 115 TALs were detected, including three within balancing-selection regions and 14 within positive-selection regions, predominantly overlapping improvement sweeps (**Figure 4A, Table S6**).

**Figure 4.**
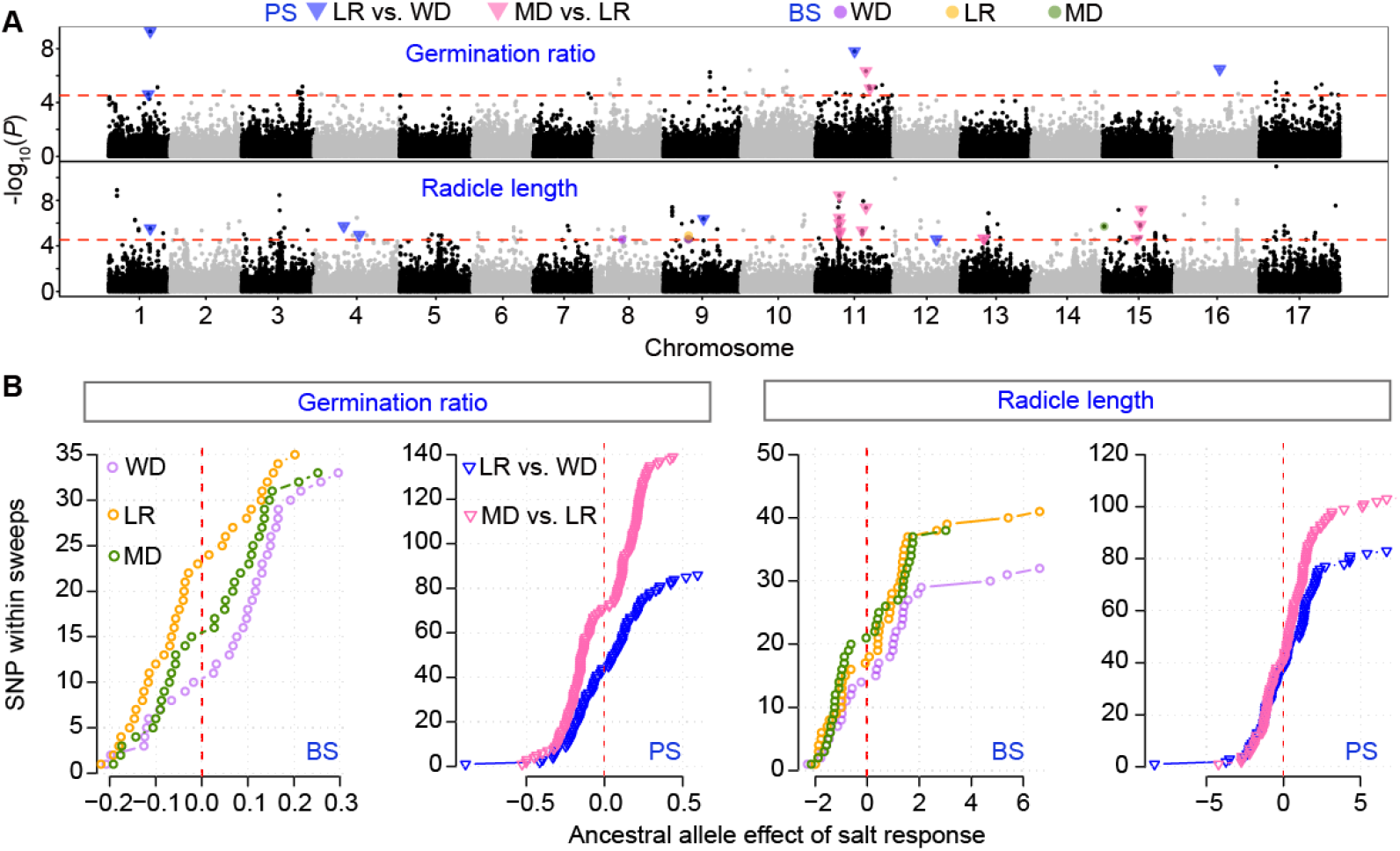
GWAS results for salt-responsive traits of germination ratio and radicle length. (**A**) Stacked Manhattan plots for salt-responsive (SR) traits. The red horizontal dashed line denotes the genome-wide significance threshold. Colored points and triangles indicate significant SNPs that overlap with regions under balancing selection (BS) or positive selection (PS). (**B**) Effects of the ancestral allele for SNPs located within BS or PS regions. Trait responsive values are shown across wild (WD), landrace (LR), and modern (MD) accessions.

We then examined the ancestral allele effect of the SNPs located within positive selection and balancing selection regions (**Figure 4B**). In general, ancestral alleles in positive selection regions showed a wider range of effects, with several SNPs exhibiting relatively large salt-response effects. In contrast, the ancestral alleles in the balancing selection regions showed a weaker salt-responsive effect. In the balanced regions, ancestral alleles tend to yield higher SR values in wild accessions than in landrace and modern groups. This suggests that, although individual SNP effects vary considerably, balanced sites contain alleles whose salt-response effects shift across populations.

## Discussion

Our findings show that sunflower evolution has been shaped by two major selective forces operating at different timescales. Ancient balancing selection preserved allelic diversity at genes related to ion transport, stress signaling, and AP2-EREBP transcriptional regulation — gene functions that likely allowed wild sunflower to cope with heterogeneous environments, consistent with previous reports of high stress-adaptive diversity in wild Helianthus (Seiler et al., 2017; Tang and Knapp, 2003). In contrast, domestication and improvement introduced strong positive selection sweeps in flowering time, oil biosynthesis, and disease-response pathways, aligning with earlier genomic studies of sunflower breeding (Badouin et al., 2017; Hübner et al., 2019). The coexistence of long-maintained polymorphisms and human-driven selective sweeps creates a mosaic genomic architecture that reflects both ecological adaptation and modern agronomic selection.

A notable consequence of these directional sweeps is the increased accumulation of deleterious mutations in cultivated sunflowers. Similar to patterns reported in maize, rice, and wheat (Fu and Horbach, 2025; Lozano et al., 2021; Lu et al., 2006), regions under strong positive selection exhibited significantly elevated genetic load, likely due to hitchhiking during rapid allele fixation and reduced efficacy of purifying selection (Yang et al., 2017). Although balanced regions also retained slightly deleterious variants, their enrichment was weaker, suggesting that long-term maintenance of polymorphism likely tolerates small-effect mutations without dramatically amplifying the genetic load.

Integrating evolutionary signatures with salt-response phenotypes revealed that adaptive stress-tolerance alleles have decreased in frequency during improvement. Ancestral alleles, particularly those maintained by long-term balancing selection, tended to confer stronger salt responsiveness in wild accessions but became rare in modern cultivars, paralleling observations in maize, sorghum, and tomato where domestication reduced abiotic-stress tolerance (Hufford et al., 2012; Lin et al., 2014; Wu et al., 2022). These findings suggest a practical breeding strategy: reintroducing or prioritizing ancestral beneficial alleles through genomic selection with advanced phenomics technology (Jin et al., 2025), targeted introgression, or genome editing (Qin et al., 2025). Our results, therefore, provide a framework for leveraging evolutionary information to guide future improvements in sunflowers under increasingly challenging environments.

## Data Availability

Supplementary data and code are available on the GitHub repository: https://github.com/GenXu1/Sunflower_domestication.

## Acknowledgements

This research was funded by the Agriculture and Food Research Initiative (grant numbers 2022-67013-36560) from the USDA National Institute of Food and Agriculture. Financial support for Yavuz Delen was provided by the Ministry of Education of the Republic of Turkey.

## Author Contributions

G.X., Y.D., I.D., and J.Y. designed this work. Y.D. and S.P.-D. conducted the cross-validation experiment. G.X., S.S.M.V.S., J.Y., and Y.D. analyzed the data. I.D. and H.W. provided conceptual advice. G.X., Y.D., and J.Y. wrote the manuscript. All authors reviewed the manuscript.

## Supporting Information

### Supplementary Tables

**Table S1**. List of accessions included in the analyses. (https://github.com/GenXu1/Sunflower_domestication/tree/main/02table/TableS1_sample_info.xlsx)

**Table S2**. Balanced loci detected in wild, landrace, and modern sunflower groups. (https://github.com/GenXu1/Sunflower_domestication/tree/main/02table/TableS2_B2_loci.xlsx)

**Table S3**. Stress-related candidate genes under balancing selection. (https://github.com/GenXu1/Sunflower_domestication/tree/main/02table/TableS3_candidate_gene_under_B2.xlsx)

**Table S4**. Positive selection loci detected during sunflower domestication and improvement. (https://github.com/GenXu1/Sunflower_domestication/tree/main/02table/TableS4_positive_selection_loci.xlsx)

**Table S5**. Candidate genes under positive selection. (https://github.com/GenXu1/Sunflower_domestication/tree/main/02table/TableS5_candidate_gene_under_positive_selection.xlsx)

**Table S6**. GWAS results for salt-responsive traits. (https://github.com/GenXu1/Sunflower_domestication/tree/main/02table/TableS6_positive_selection_loci.xlsx)

## Supplementary Figures

**Figure S1.**
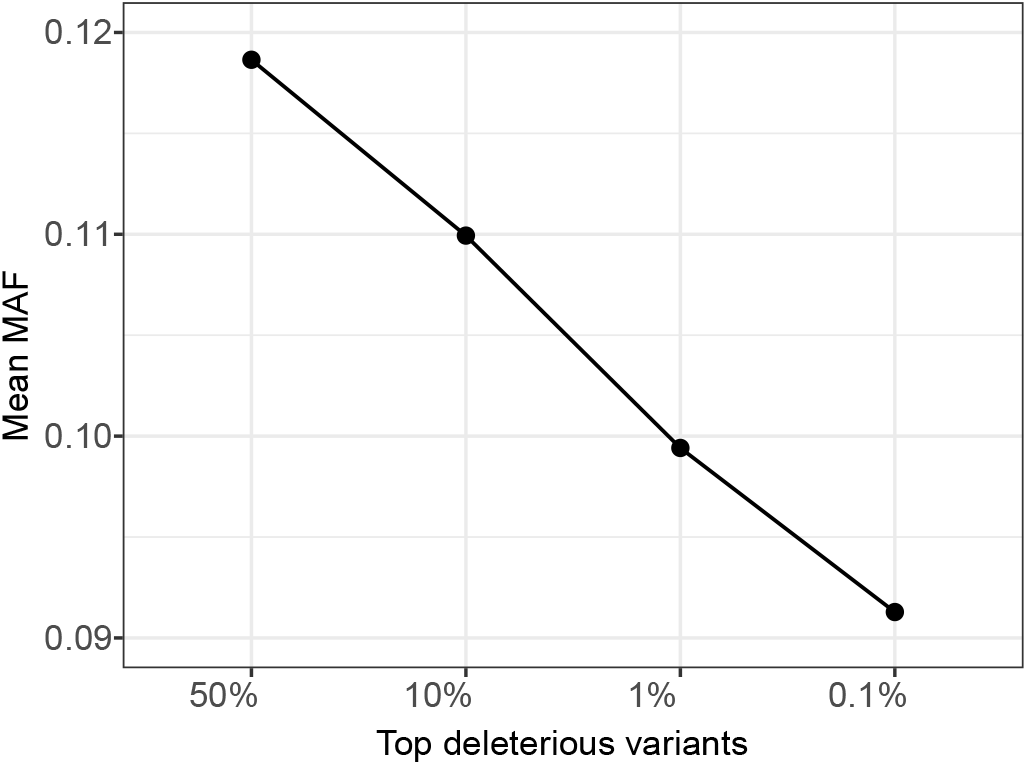
Mean minor allele frequency (MAF) of deleterious variants across different deleteriousness thresholds.

**Figure S2.**
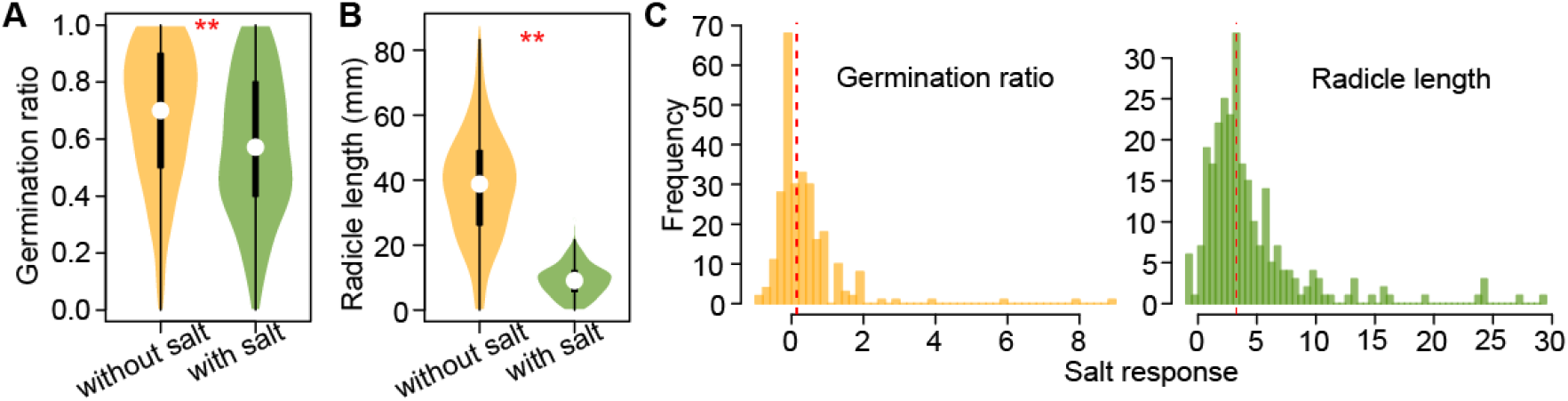
Germination ratio and radicle length under different salt conditions. (**A-B**) Distribution of germination ratio (A) and radicle length (B) under salinity (with salt) and the control group (without salt). Asterisks indicate significance levels based on paired Wilcoxon’s test (**, *P* ≤ 0.01). (**C**) The distributions of salt-responsive (SR) traits. The red dashed lines denote the median values.

